# Tim29 is required for stem cell activity during regeneration in the flatworm *Macrostomum lignano*

**DOI:** 10.1101/2020.05.29.123885

**Authors:** Stijn Mouton, Kirill Ustyantsev, Frank Beltman, Lisa Glazenburg, Eugene Berezikov

## Abstract

Domain of Unknown Function (DUF) genes are broadly conserved in metazoa but have an unknown biological function. Previously, we established transcriptional profiles of the germline and stem cells of the flatworm *Macrostomum lignano* (Grudniewska et al., 2016). Here, we used this information to probe the function of DUF genes in *M. lignano*. Recently, Kang et al. (2016) found that DUF2366 is a mitochondrial inner membrane protein that interacts with the protein import complex TIM22. DUF2366 was renamed TIM29, and shown to stabilize the TIM22 complex, but its biological role remained largely unknown. We now demonstrate that DUF2366/TIM29 is dispensable in the homeostatic condition in *M. lignano*, but is essential to adapt to a highly proliferative state required for regeneration. Our results show that ‘de-DUFing’ the function of DUF genes might require special biological conditions, such as regeneration, and establishes *M. lignano* as an informative platform to study DUF genes.

## Introduction

Protein domains of unknown function (DUFs) are often neglected, although they represent a treasure trove of unknown biology (Bateman et al., 2010; Jaroszewski et al., 2009). Domains represent the functional units of proteins and typically have distinct structures and functions. Despite decades of research, more than 20% of all domains in the Pfam database, the so-called DUFs, are still functionally uncharacterized (Bateman et al., 2010; Goodacre et al., 2013; Mudgal et al., 2015). Evolutionary conservation suggests that many of these DUFs are important, but studies indicated that they more likely represent biological functions specific to certain conditions, or certain groups of organisms, rather than being part of the core machinery common to all life (Bateman et al., 2010; Goodacre et al., 2013). This does not reduce their value, as their undiscovered functionality can represent novel biochemical pathways, alternative solutions to known reactions, or new regulatory mechanisms (Jaroszewski et al., 2009). In the context of stem cell biology and regeneration, investigating DUFs can result in identifying novel aspects of the *in vivo* regulation of stem cells, which could provide unexpected breakthroughs for both fundamental and biomedical research. To study *in vivo* stem cell biology during development, adult tissue turnover, regeneration, and ageing, various model organisms are used. An increasingly attractive model is the free-living hermaphrodite flatworm *Macrostomum lignano* (Wudarski et al., 2020). *M. lignano* is a transparent worm with a large mesodermal population of proliferating neoblasts, which represent flatworm stem cells and progenitors (Ladurner et al., 2008, 2000). These neoblasts enable a high cellular turnover during adult homeostasis (Nimeth et al., 2002) and a large regeneration capacity (Egger et al., 2006; Nimeth et al., 2007). After amputation or incision, *M. lignano* can regenerate any part posterior of the pharynx and the anterior-most body part (the rostrum), although a head cannot be regenerated (Egger et al., 2006). Molecular research of the neoblasts was facilitated by the development of multiple methods, such as *in situ* hybridization, RNA interference, and transgenesis (Grudniewska et al., 2016; Pfister et al., 2008, 2007; Wudarski et al., 2017), and the assembly of a high quality genome and transcriptome (Grudniewska et al., 2018; Wudarski et al., 2017). In 2016, we established transcriptional signatures of proliferating somatic neoblasts and germline cells by performing RNA-seq of FACS-isolated cells of worms in different conditions (Grudniewska et al., 2016). This dataset represents a convenient resource to identify DUFs with functions related to *in vivo* stem cell and germline regulation, regeneration, and development. It is expected that many of these regulators are conserved between flatworms and human, since about 47% of all *M. lignano* transcripts have human homologs, while for neoblast-enriched transcripts this is even about 85% (Grudniewska et al., 2018).

In this paper, we focus on one example: DUF2366. According to the Pfam database, this family of proteins is widely conserved from nematodes to humans. During our characterization of DUF2366 in *M. lignano*, two manuscripts were published, which identified DUF2366 (named C19orf53 in human) as a novel subunit of the human Translocase of the Inner Membrane 22 (TIM22) complex in HEK cells (Callegari et al., 2016; Kang et al., 2016). It was demonstrated that DUF2366 is required for maintaining the structural integrity and the assembly of the TIM22 complex, which mediates the import and insertion of hydrophobic proteins into the mitochondrial inner membrane (Callegari et al., 2016; Kang et al., 2016). Consequently, DUF2366 was renamed TIM29 (Callegari et al., 2016; Kang et al., 2016). In addition, it was suggested that TIM29 contacts the Translocase of the Outer Membrane (TOM) complex, enabling transport of hydrophobic carrier substrates across the aqueous intermembrane space (Kang et al., 2016). Interestingly, both papers studied the effect of TIM29 RNA interference (RNAi) on HEK cell proliferation, and reported contradicting results. While Kang et al. did not observe a significant effect of hTIM22 knockdown on cell proliferation (Kang et al., 2016), Callegari et al. observed a significantly decreased cell proliferation (Callegari et al., 2016). In other words, the importance of DUF2366/TIM29 for stem cell biology remained unclear.

Here, we identify the DUFs conserved in *Macrostomum lignano* and demonstrate the crucial role of Mlig-DUF2366/TIM29 for *in vivo* stem cell activity during whole body regeneration by means of RNA interference studies.

## Results

### Identification of uncharacterized proteins in *M. lignano*

To facilitate the discovery of novel genes involved in stem cell function, germline biology, regeneration, and development, we identified all genes in the *M. lignano* genome-guided transcriptome assembly Mlig_3_7_DV1_v3 (Grudniewska et al., 2018) encoding uncharacterized proteins (Suppl. Table 1). Due to partial genome duplication and redundancy, very closely related genes were grouped using Corset (Davidson and Oshlack, 2014) into so-called transcript clusters for the downstream analysis (Grudniewska et al., 2018; Wudarski et al., 2017). Of the 820 identified DUF transcript clusters, 274 have identifiable homologs in human. Based on the expression level of the DUFs in different conditions and using previously established neoblast and germline transcriptional signatures (Grudniewska et al., 2016), categories were provided to predict their functional role in stem cells, the germline, regeneration, and development (Table 1). The value of this candidate list was tested with a pilot RNA interference (RNAi) screen of three randomly chosen genes coding uncharacterized proteins: *DUF2315* (Mlig002791.g5), *UPF0197* (Mlig006314.g7), and *DUF2366* (Mlig032364.g1). The screen focused on triple tail regeneration of *M. lignano* (Figure 1A), which depends on functional neoblasts. The decision to repeatedly amputate the tail was based on the experience obtained during previous RNAi screens indicating that some genes require a longer knockdown treatment than others to result in an obvious phenotype (Grudniewska et al., 2016). Knockdown of one of the three genes, *Mlig-DUF2366*, resulted in a reproducible phenotype. After 28 days, all *DUF2366*(RNAi) worms failed to regenerate new tissue, while all *gfp*(RNAi) worms, representing the negative control, successfully regenerated the tail (Figure 1C). Interestingly, at least three repeated amputations of the tail-plate are necessary to induce this phenotype in 100% of the *DUF2366*(RNAi) worms. After a single tail-amputation, all *DUF2366*(RNAi) worms were still able to regenerate a tail (Figure 1B), and two tail-amputations demonstrated a variable degree of regeneration between *DUF2366*(RNAi) worms. Taken together, this suggests that without *Mlig-DUF2366* expression, worms have limited regenerative abilities. Based on these results, we decided to further characterize *DUF2366* in *M. lignano*, focusing on its requirement for stem cell function.

**Table 1.** Categories of DUF genes in *M. lignano*

| Category | Transcript clusters* | DUFs |
| --- | --- | --- |
| Neoblasts | 13 | 11 |
| Germline | 36 | 25 |
| Regeneration | 126 | 78 |
| Development | 289 | 168 |
| Unknown | 425 | 198 |

\* same transcript cluster can belong to several categories.

**Figure 1.**
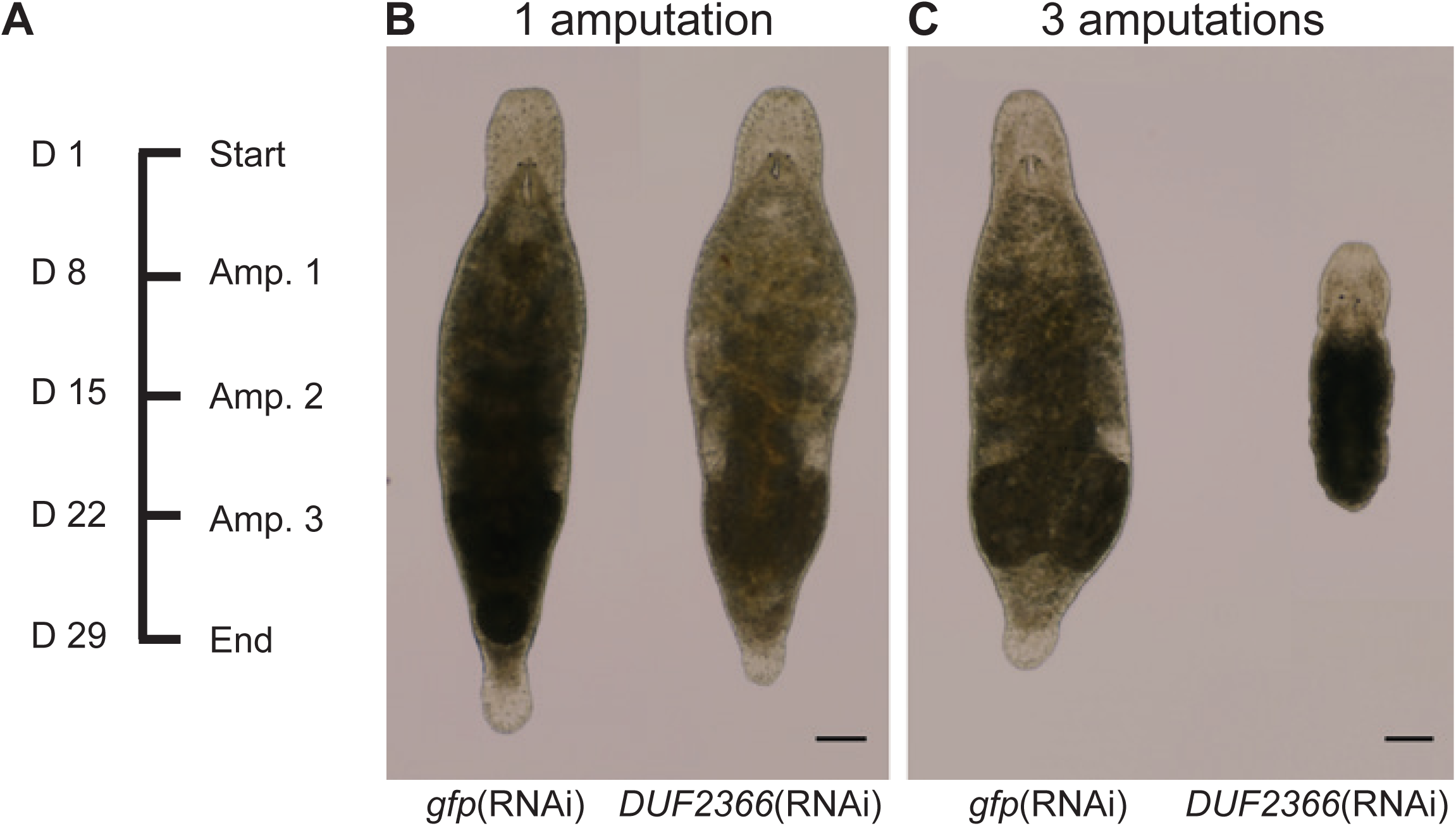
RNA interference screen. (**A**) Experimental design. D represents the time of treatment in days. Amp. describes the number of tail-amputations. (**B**) After a single amputation, worms are able to regenerate the tail within a week. (**C**) After the third amputation of the tail, *gfp*(RNAi) worms can regenerate, while *DUF2366*(RNAi) worms show a complete lack of regeneration. Scale bars are 100 µm.

### *M. lignano DUF2366* has an enriched expression in neoblasts and is homologous to TIM29

The *Mlig-DUF2366* gene has three nearly identical loci in the Mlig_3_7_DV1 genome assembly (Grudniewska et al., 2018): Mlig032364.g1, Mlig015320.g2, and Mlig18840.g2. All three *Mlig-DUF2366* loci have strong homology to the TIM29 protein superfamily members (Pfam, PF10171) from diverse Metazoa (Suppl. Figure 1).

The previously obtained transcriptional profiles of sorted cells (Grudniewska et al., 2016) demonstrated that *DUF2366* transcripts have an elevated expression in proliferating somatic neoblasts compared to differentiated cells (Suppl. Table 1). Interestingly, according to the online PlanMine resource (Brandl et al., 2016), the *Schmidtea mediterranea* homolog, dd_Smed_v6_9413_0_1 (Suppl. Figure 1), also has a higher expression in X1 cells (cycling stem cells), compared to X2 (progenitors) and Xins cells (differentiated cells), and is included in the ‘Stem cells versus differentiated cells_low stringency’-list. This suggests that enriched expression of *DUF2366* in neoblasts is conserved in multiple flatworm species.

Based on the two recent papers renaming the human DUF2366 as TIM29 (Callegari et al., 2016; Kang et al., 2016), a mitochondrial localization of the Mlig-DUF2366 protein can be expected. Indeed, the online MitoFates tool (Fukasawa et al., 2015) predicts that the translated protein sequence of *DUF2366* transcripts contain a mitochondrial presequence and TOM20 recognition motifs. MitoFates also predicts a mitochondrial presequence for dd_Smed_v6_9413_0_1 (PlanMine) and human TIM29 (GeneBank accession: NP_612367.1) translated protein sequences (data not shown).

### Experimental setup to study the role of *Mlig-DUF2366/TIM29* in neoblasts and regeneration

To investigate the potential stem cell function of DUF2366, a set of RNAi experiments was performed using *M. lignano*. In total, four specific experimental ‘classes’ were characterized: *gfp*(RNAi) uncut, *gfp*(RNAi) cut, *Mlig-DUF2366*(RNAi) uncut, and *Mlig-DUF2366*(RNAi) cut (Figure 2A). As GFP is not expressed in wild type worms, the *gfp*(RNAi) classes represent the negative control. The uncut conditions represent worms in which proliferation is only required for cell turnover during adult tissue homeostasis, and the production of gametes. In the cut conditions, the body was amputated by cutting worms between the pharynx and testes after 1 week of RNAi treatment, inducing regeneration of the whole body. Flatworm regeneration is a convenient readout for stem cell functionality, as it requires neoblast proliferation, migration, and differentiation. Whole-body regeneration was chosen as a specific condition potentially requiring DUF2366 activity, based on the results of the preliminary screen, and the notion that the important function of DUFs is often restricted to specific conditions.

**Figure 2.** ***Mlig-****DUF2366*(RNAi) study. (**A**) Experimental design. The horizontal grey squares indicate which worms are used at each time point, represented by the number of days. **(B)** After 4 weeks of RNAi treatment, most *Mlig-DUF2366*(RNAi) worms look similar to the *gfp*(RNAi) controls, although a few individuals demonstrate a limited capacity of tissue turnover. This is illustrated by the worm on the right which shrank, degenerated the gonads, and developed bulges in the epidermis. (**C**) The effect of RNA interference on the number of mitotic cells. *Mlig-DUF2366*(RNAi) causes a significant decrease in the number of mitotic cells in regenerating worms (Regeneration), but not in uncut worms (Homeostasis whole body). To allow comparison between cut and uncut worms, the number of mitotic cells was quantified in the head and pharynx region of uncut worms (homeostasis anterior of testes) which correspond with the fragment left after amputating the body. In control *gfp*(RNAi) worms, regeneration causes a significant increase in the number of mitotic cells. This is not observed in *Mlig-DUF2366*(RNAi) worms. n.s.: p > 0.05, ***p < 0.001 (ANOVA and post-hoc Tukey test). Genes differentially expressed between uncut *Mlig-DUF2366*(RNAi) and *gfp*(RNAi) worms. (**D**) Genes differentially expressed between *gfp*(RNAi) and *Mlig-DUF2366*(RNAi) worms. (**E**) During knockdown of *DUF2366*, worms are not able to regenerate the body. Post Amp. represents time after amputation of the body. All scalebars are 100 µm.

Different measurements were performed on all four RNAi classes (Figure 2A). First, the morphology of all four classes and the regeneration capacity of cut worms were assessed. Second, the number of mitotic cells was quantified, which serves as a marker for proliferation rate. Third, gene expression was studied by means of RNA sequencing (Suppl. Table 2). Both mitotic labeling and RNA-seq were performed on the tenth day of RNAi, which represents the second day of regeneration in the cut worms. Fourth, to interpret the changes in gene expression, Gene Ontology (GO) Term analysis of differentially expressed genes was performed.

### *Mlig-DUF2366/TIM29* knockdown during homeostasis does not result in a prominent phenotype

After four weeks of RNAi, the majority of uncut *Mlig-DUF2366*(RNAi) worms still had a similar morphology as the uncut *gfp*(RNAi) worms (Figure 2B). A limited number of individuals, however, showed an impaired maintenance of the body by e.g. degeneration of the gonads, degeneration of the rostrum, a shrinking size, and appearance of small bulges (Figure 2B), which often resulted in death of the individual. At the tenth day of RNAi, the number of mitotic cells, representing dividing somatic neoblasts and germline cells, was not significantly different between homeostatic control and *Mlig-DUF2366*(RNAi) worms (p=0.610, ANOVA and post-hoc Tukey test) (Figure 2C). This demonstrates that *Mlig-DUF2366*(RNAi) worms can maintain the proliferation rate required for cellular turnover during homeostasis. Differential gene expression analysis identified only 23 significantly differentially expressed genes when uncut *Mlig-DUF2366*(RNAi) and uncut *gfp*(RNAi) worms were compared (Figure 2D). In other words, both groups of worms only show minimal differences at the molecular level. In conclusion, knockdown of *Mlig-DUF2366* in uncut worms did not result in a consistent prominent phenotype within the timeframe of four weeks.

### *Mlig-DUF2366/TIM29* knockdown impairs whole-body regeneration

The regenerative ability was assessed at one and two weeks after amputation of the body, and was clearly inhibited by knockdown of *Mlig-DUF2366* at both time points. One week after amputation, all *Mlig-DUF2366*(RNAi) worms had a smaller tail than the negative controls, although a large variation could be observed between different individuals. In worst case, there was a complete lack of regeneration, while in best case worms regenerated a small tail plate (Figure 2E). With advancing time, the phenotype became more apparent. Two weeks after amputation, the *gfp*(RNAi) worms were completely regenerated and resembled young adults. In contrast, the *Mlig-DUF2366*(RNAi) worms were small and disproportionate. Due to the impaired regenerative tissue remodeling, the location of amputation could still be observed. The morphology of the tail still varied from lacking to a small tail plate. In several cases, the appearance of bulges could be observed (Figure 2E).

### *Mlig-DUF2366/TIM29* is required for adapting to regenerative conditions

The observation that knockdown of *Mlig-DUF2366* results in a consistent phenotype only during triple tail-regeneration and whole-body regeneration indicates that worms cannot adapt to highly proliferative conditions during RNAi treatment. To understand the underlying mechanisms, it is important to compare changes induced by whole-body regeneration in *Mlig-DUF2366*(RNAi) and *gfp*(RNAi) animals.

Quantification of mitotic neoblasts in the amputated head-fragments and the corresponding region of the body, anterior of the testes, in uncut worms presents the first confirmation for this inability to adapt. In *gfp*(RNAi) worms, regeneration causes a significant increase in the number of mitotic neoblasts, demonstrating increased proliferation during regeneration (p=0.001, ANOVA and post-hoc Tukey test) (Figure 2C). During *Mlig-DUF2366* knockdown, inducing regeneration does not significantly increase the number of mitotic neoblasts (p=0.999, ANOVA and post-hoc Tukey test) (Figure 2C).

The inhibited adaptation to regenerative conditions due to *Mlig-DUF2366* knockdown can also be observed at the level of gene expression, and 3854 significantly differentially expressed genes were identified when comparing cut *Mlig-DUF2366*(RNAi) and cut *gfp*(RNAi) worms, of which 1827 were upregulated and 2027 were downregulated (Figure 2D). In *gfp*(RNAi) worms, genes with an upregulated expression due to regeneration are enriched for neoblast transcripts (1.37-fold; p < 10^−12^, Pearson’s Chi-squared test with Yates’ continuity correction), while in *Mlig-DUF2366*(RNAi) worms neoblast transcripts are depleted among genes with increased expression due to regeneration (0.44-fold, p < 10^−12^) (Suppl. Table 3).

### Knockdown of *Mlig-DUF2366/TIM29* has a global cellular impact in regenerating worms

To better understand the impaired regenerative response during *Mlig-DUF2366*(RNAi), we compared gene expression between cut *Mlig-DUF2366*(RNAi) and cut *gfp*(RNAi) worms in more detail by means of GO term analysis. This analysis shows that multiple molecular biological processes are affected due to knockdown of *Mlig-DUF2366*, including translation and protein transport, metabolism, transcription regulation and RNA processing, cell division, and mitochondrial function (Figure 3). Interestingly, biological clusters of significantly enriched GO terms are only found for downregulated genes. Thus, knockdown of *Mlig-DUF2366* has a global effect, impacting multiple basic molecular processes and organelles in regenerating worms.

**Figure 3.**
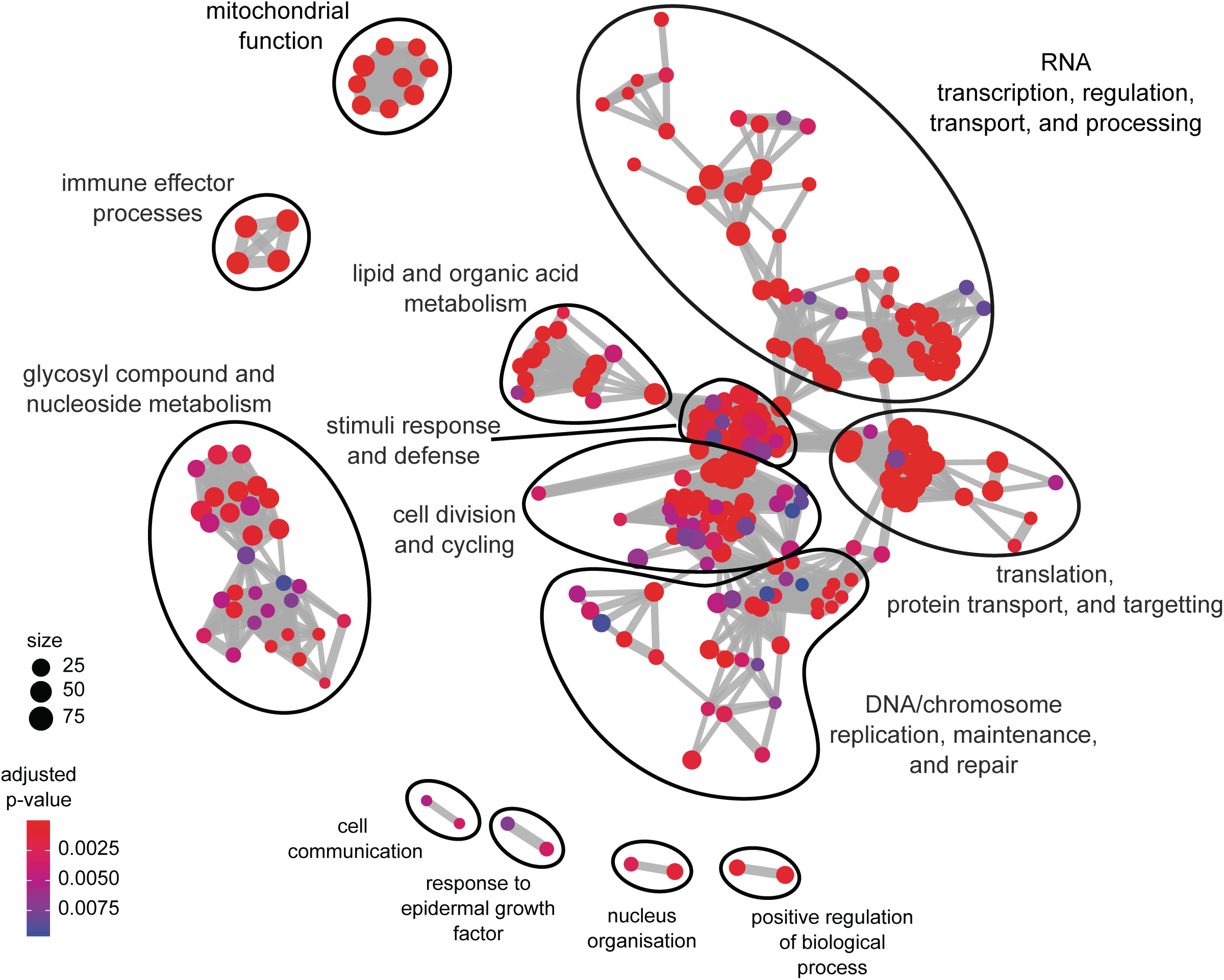
Enrichment of GO terms of biological processes categories downregulated in cut *Mlig-DUF2366*(RNAi) vs cut *gfp*(RNAi) worms represented as a graph. Each GO term is shown as a circle, a node in the graph, which size is proportional to the number of differentially expressed *M. lignano* human homologues assigned to the term, and the nodes color corresponds to adjusted p-values supporting its enrichment. Semantically related GO term nodes are connected with an edge. Closely related GO terms are arbitrarily outlined and given generalized names, based on the most frequent GO terms in each group.

## Discussion

All data together demonstrates that knockdown of one gene, *Mlig-DUF2366*, is enough to impair whole-body regeneration by inhibiting the adaptation to highly proliferative conditions. In other words, *Mlig-DUF2366* is identified as an essential gene for stem cell function. Interestingly, computational analysis indicated a mitochondrial localization of the DUF2366 protein in *M. lignano*, which is consistent with the recent renaming of the human DUF2366 as TIM29 based on its characterization as an inner mitochondrial membrane protein in HEK cells (Callegari et al., 2016; Kang et al., 2016). This mitochondrial localization and function explains the enrichment of several GO term processes and components related to mitochondria in the differentially expressed genes between cut *Mlig-DUF2366*(RNAi) and cut *gfp*(RNAi) worms (Figure 3). Moreover, the described function of human TIM29 provides a framework to explain how knockdown of *Mlig-DUF2366* can cause the inability to adapt to highly proliferative conditions, and consequently limit the regenerative capacity (Figure 4). TIM29 is required for stabilizing the structural integrity and the assembly of the TIM22 complex, an important protein import machinery in the inner mitochondrial membrane. Consequently, a lack of TIM29 will result in a failing translocation and integration of proteins, such as metabolite carrier proteins and subunits of the TIM22 and TIM23 complex, into the inner mitochondrial membrane (Callegari et al., 2016; Kang et al., 2016). This will impact energy production, mitochondrial proteostasis, and cause perturbations of mitochondrial physiology, which are all potential causes of mitochondrial stress (Durieux et al., 2011; Houtkooper et al., 2013; Jensen and Jasper, 2014; Samluk et al., 2018; Yoneda et al., 2004). Mitochondrial stress can trigger a vigorous transcriptional response aimed at stabilizing mitochondrial and cellular functions, promoting cellular repair, and adapting the metabolism to stress conditions (Jensen and Jasper, 2014; Lin and Haynes, 2016). The observed global changes in metabolism, translation, transcription regulation, DNA repair, stimuli response, and mitochondrial function support this hypothesis (Figure 3). This state of stress inhibits the required molecular response for successful regeneration and growth. As a result, worms are not able to increase the proliferation rate required for whole-body regeneration, which is shown at both the cellular and molecular level (Figure 2).

**Figure 4.**
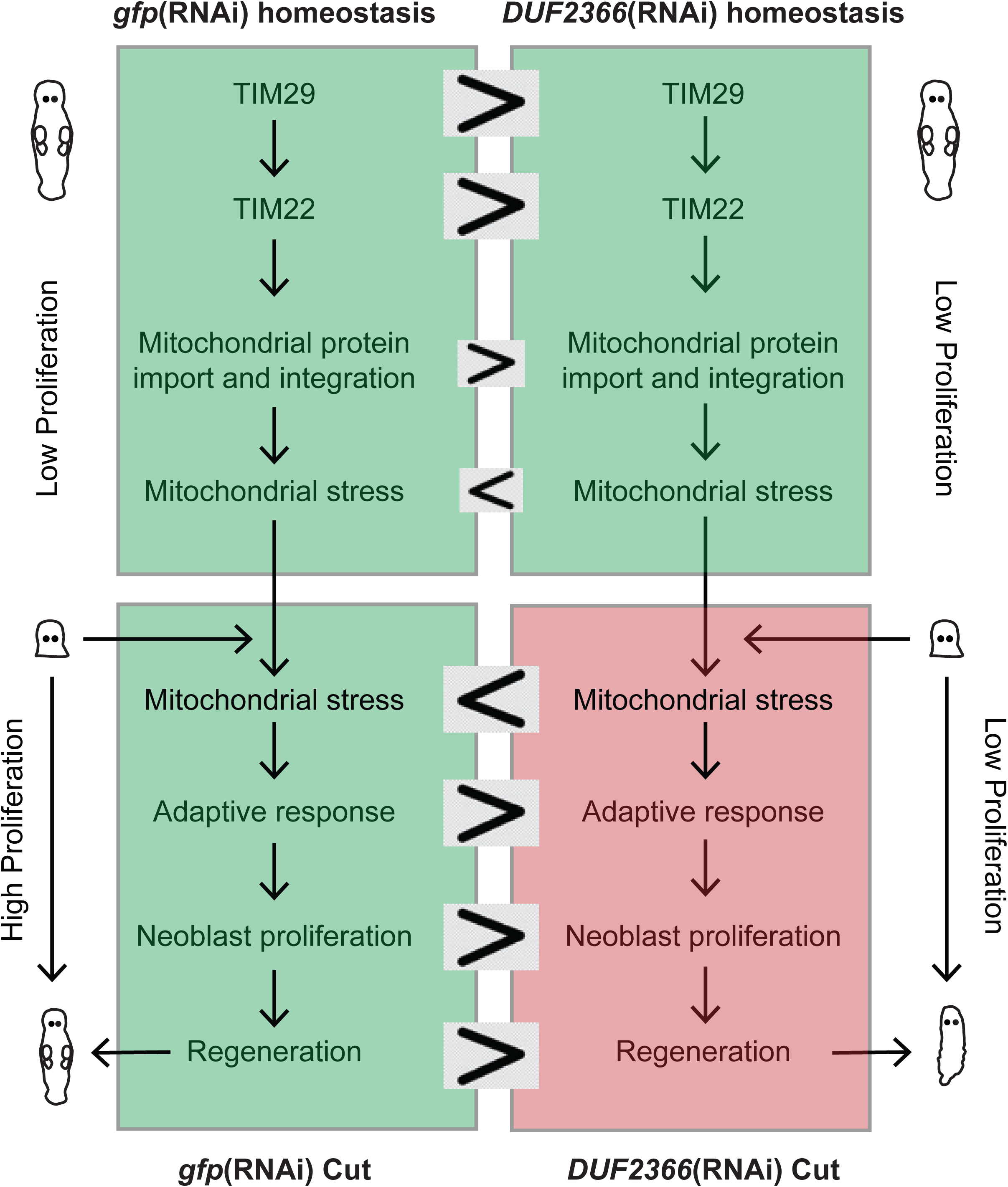
Schematic visualization explaining how knockdown of *Mlig-DUF2366/TIM29* results in a regenerative phenotype.

Our findings demonstrate that healthy mitochondria are essential for proper neoblast function, and add to the increasing recognition of mitochondrial signaling as a key component to mediate stem cell activity. The emerging picture is that mitochondria continuously integrate cellular and environmental cues to influence stem cell fate and activity, which enables organisms to adapt to the environmental changes (Battersby and Richter, 2013; Lisowski et al., 2018; Zhang et al., 2018). Many questions remain, however, as the majority of published mitochondrial research focused on post-mitotic tissues, and the role of mitochondria in the context of stem cells has been largely neglected until recently for several reasons (Zhang et al., 2018). First, stem cells mainly rely on glycolysis for their energy, although metabolic properties differ between different types of stem cells. Second, many stem cells have a low abundance of mitochondria (Rafalski et al., n.d.; Zhang et al., 2018). Similarly, neoblasts are described as small spherical cells with only a thin rim of cytoplasm containing free ribosomes and mitochondria. In stages which were assumed to be more differentiated, also rough endoplasmic reticulum and Golgi complex were sporadically observed (Bode et al., 2006; Rieger et al., 1999). Despite the low abundance of cytoplasmic organelles in neoblasts, we selected *Mlig-DUF2366* based on its enriched expression in proliferating neoblasts. A finding which is confirmed in the planarian *Schmidtea mediterranea*, based on the gene expression analysis presented in the online PlanMine (Brandl et al., 2016) tool. It is unknown why an inner mitochondrial membrane protein would have an enriched expression in the neoblasts and why the variation in expression of *Mlig-DUF2366* varies more in samples of proliferating cells than in G1 cells. We suggest that the elevated expression could be linked to the proliferation rate of the neoblasts, as increased production of new cells could require increased levels of mitochondrial biogenesis (Figure 4). This reasoning can also explain why a threshold of regeneration is needed to induce a consistent phenotype, e.g. the need for at least three amputation of the tail to obtain an obvious regeneration impairment in 100% of the worms. If increased production of new cells boosts mitochondrial biogenesis, it will also enhance mitochondrial stress and increase the number of cells experiencing this stress in the absence of DUF2366 during RNAi treatment. From this point of view, variation in the level of proliferation due to a combined effect of e.g. growth and unintended injury by transferring, could explain why some uncut worms show a failing cellular turnover during knockdown of *Mlig-DUF2366*, while the majority does not.

The development of a *Mlig-DUF2366*(RNAi)-phenotype only during large scale regeneration fits with the description of DUF proteins having an important biological role limited to specific conditions (Bateman et al., 2010; Goodacre et al., 2013). It is therefore essential to study the function of DUFs in different conditions, and this manuscript demonstrates the value of *Macrostomum lignano* for this type of research. *M. lignano* provides an *in vivo* system enabling to study stem cells in their natural environment. Different conditions besides cellular turnover during adult homeostasis can be easily induced. Examples are different levels of regeneration by amputating different portions of the body, development, starvation, growth and even degrowth based on the available amount of food. Moreover, the expanding molecular toolbox and especially the recently developed methods of transgenesis will further facilitate *in vivo* studies to identify and characterize a stem cell function of Uncharacterized Proteins (Mouton et al., 2018; Wudarski et al., 2017). The here presented list of Uncharacterized Proteins presents an ideal starting point for selecting candidates. The value of this list is clearly demonstrated as screening three candidates was sufficient for identifying a candidate which is essential for neoblast activity. The fact that this is a mitochondrial protein and not a direct cell cycle regulator only adds to the value, as screening more candidates will provide novel insight into which processes and organelles are essential for stem cell biology.

## Materials and methods

### Culture of *Macrostomum lignano*

The free-living flatworm *M. lignano* is cultured in Petri dishes with nutrient-enriched artificial seawater (f/2), at a temperature of 20°C and a 14h/10h light/dark cycle (Anderson et al., 2005). Worms are fed *ad libitum* with the diatom *Nitzschia curvilineata* (Rieger et al., 1988).

### Homology detection and sequence analysis

To find a homology to other known protein families, nucleotide sequences of the transcribed *M. lignano* DUF2366 loci as well as *S. mediterranea* dd_Smed_v6_9413_0_1 transcript sequence (PlanMine) were directly submitted to the NCBI Conserved Domain Search server (Lu et al., 2020). Open reading frames (ORFs) analysis, multiple sequence alignments construction and visualization were done in Uinpro UGENE v34.0 (Okonechnikov et al., 2012). Amino acid sequences of TIM29 conserved domain family of other Metazoa species were obtained directly from Pfam (https://pfam.xfam.org/family/PF10171#). Prediction of mitochondrial processing presequence and TOM20 recognition motifs was performed using the online MitoFates tool (Fukasawa et al., 2015) submitting translated ORFs sequences.

### RNA interferenc

The production of dsRNA was performed following the published protocol (Grudniewska et al., 2018, 2016), and the used primers of candidate genes are presented in (Suppl. Table 4). Candidate genes were knocked down by means of RNAi with double-stranded RNA delivered by soaking as previously described (De Mulder et al., 2009; Grudniewska et al., 2016). The RNAi soaking experiments were performed in 24-well plates in which diatoms were grown. Individual wells contained 300 µl of dsRNA solution (10 ng/µl in f/2 medium) in which 15 individuals were maintained. The preliminary RNAi-screen lasted for 4 weeks, and the tail of worms was amputated after 1, 2, and 3 weeks of RNAi treatment. For the *Mlig-DUF2366*(RNAi)-screen, worms of the regenerative condition were treated for 3 weeks, and the body was amputated by cutting the worms in the region between the testes and the pharynx after 1 week of RNAi treatment. Worms of the homeostasis condition were treated for 4 weeks. In all RNAi treatments, animals were weekly transferred to fresh 24-well plates to ensure sufficient amounts of food. As a negative control, *gfp* dsRNA was used. To illustrate the observed changes in morphology and regeneration capacity, photos were taken using an EVOS XL Core Imaging System (ThermoFisher). For this, worms were temporary relaxed in 1:1 f/2:MgCl_2_.6H_2_O (7.14%) in a small drop in a Petri dish.

### Mitotic labeling

For both *Mlig-DUF2366*(RNAi) and *gfp*(RNAi), cut and uncut worm were collected at the tenth day of RNAi treatment. In the cut condition, this time point represents 2 days after amputation. Mitotic labeling was performed as described before (Grudniewska et al., 2016; Ladurner et al., 2000). In short, worms were washed in f/2 medium, relaxed in 1:1 f/2:MgCl_2_.6H_2_O (7.14%), and fixed in 4% paraformaldehyde (PFA) for 1h at room temperature (RT). Afterwards, they were washed with PBS-T (PBS and 0.1% Triton X-100) and blocked with BSA-T (1% bovine serum albumin in PBS-T) for 30 minutes at RT. The primary anti-phospho histone H3 Antibody (Millipore) was diluted 1:250 in BSA-T and applied overnight at 4°C, followed by washing with PBS-T at RT. Worms were then incubated in secondary goat anti-rabbit IgG Antibody conjugated with FITC (Millipore) which is diluted 1:150 in BSA-T, for 1h at RT. After being washed with PBS-T, slides were mounted using Vectashield (Vector Laboratories US, Burlingame, CA). Mitotic cells were visualized using a Leica TCS SP8 confocal microscope and counted with the Cell counter plugin in ImageJ. For each of the four conditions, the number of mitotic cells was quantified for a total of 12 individuals obtained from two independent labeling-experiments (n=2*6). To determine if the number of cells was significantly different between conditions, an ANOVA and post-hoc Tukey test were performed in SPSS.

### Preparation and sequencing of RNA-seq libraries

Worms of the four different conditions (*Mlig-DUF2366*(RNAi) uncut; *Mlig-DUF2366*(RNAi) cut; *gfp*(RNAi) uncut; *gfp*(RNAi) cut) were collected at the tenth day of RNAi treatment. In the cut condition, this time point represents 2 days after amputation. For each condition, four replicates of 45 individuals each were rinsed with f/2 medium, suspended in 500 µl TRIzol reagent (Ambion) and stored at −80°C.

RNA was extracted from the samples with the Direct-zol RNA MiniPrep Kit (Zymo Research), following the manufacturer’s protocol. RNA-Seq libraries were made using the CEL-Seq2 protocol (Hashimshony et al., 2016, 2012), and as a first step a mix of RNA, primer, spike-in, and dNTPs was made. While this method was originally designed for single cells, it also works well with larger amounts of RNA. Sequencing was performed using the T-fill protocol (Wilkening et al., 2013) on an Illumina HiSeq 2500 machine.

### Differential expression analysis of RNA-Seq data

Illumina reads were mapped to the *M. lignano* genome assembly Mlig_3_7 (Wudarski et al., 2017) using STAR software v. 2.6.0c (Dobin et al., 2013) and the transcriptome annotation version Mlig_RNA_3_7_v3 (Grudniewska et al., 2018). The transcriptome quantification option of STAR was used to derive initial transcript counts, which were consolidated into transcript cluster counts using Corset (Davidson and Oshlack, 2014). Differential gene expression analysis was performed using generalized linear models implemented in the edgeR software package (McCarthy et al., 2012). Only transcript clusters that had at least 1 count per million in at least 3 samples were included in the analysis. An FDR cutoff of 0.05 was used to establish statistically significant differentially expressed genes.

### Go Term analysis

For genes differentially expressed between cut *Mlig-DUF2366*(RNAi) vs cut *gfp*(RNAi) worms, human homolog gene annotations were extracted from the previously annotated trascriptome Mlig_RNA_3_7_v3 (Grudniewska et al., 2018). The resulted list of *M. lignano* human homologues was then processed and analyzed using a custom script written in R programming language. Libraries “org.Hs.eg.db” (Carlson, 2019) and “clusterProfiler” (Yu et al., 2012) were used for the GO term analysis, applying the function *enrichGO* with the following parameters: [List of Human ENTREZ gene IDs], OrgDb=org.Hs.eg.db, ont=“BP”, pvalueCutoff = 0.01, qvalueCutoff = 0.01, pAdjustMethod = “BH”. The results of the analysis were visualized as a graph using the *emapplot* function, showing all categories and coloring nodes by their adjusted p-values. The graph was exported to PDF and manually processed in Inkscape v.0.92 vector graphics software to apply additional annotations.

### Data accessibility

RNA-seq data have been deposited at DDBJ/EMBL/GenBank under the BioProject accession number PRJNA606131.

## Acknowledgements

The RNAi screen and characterization of the *DUF2366* knockdown phenotype was supported by the European Research Council Starting Grant (MacModel, grant no. 310765) to EB. Work on gene expression analysis performed in the Institute of Cytology and Genetics SB RAS was supported by the Russian Science Foundation grant no. 20-14-00147 to EB.

---

**Supplementary Figure 1.**
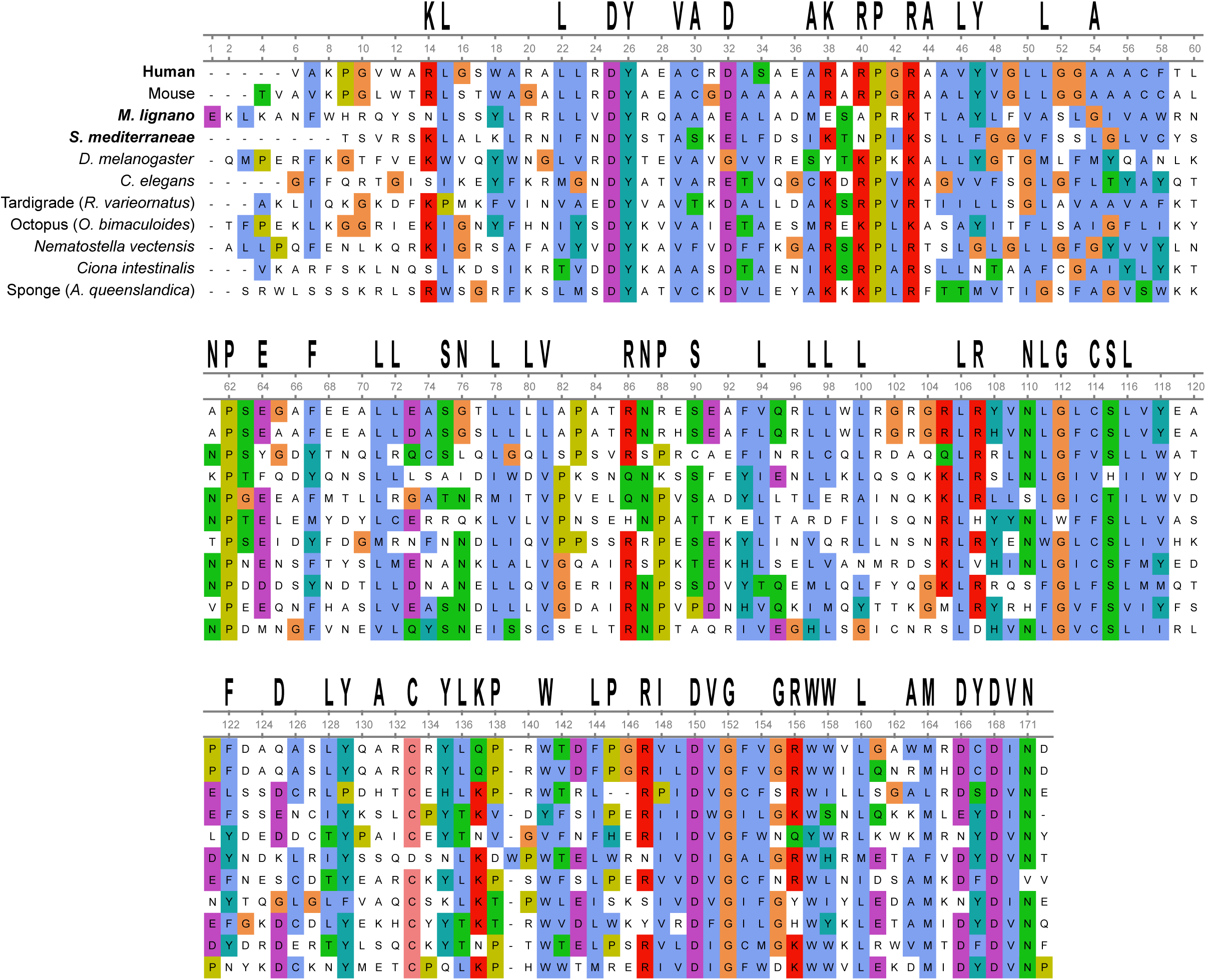
Multiple amino acid sequence alignment of TIM29 family protein domains (Pfam, PF10171) from various Metazoa species. Conserved (> 50%) amino acids are designated at the top of the alignment. *M. lignano* transcript number - Mlig018840.g2. *S. mediterranea* PlanMine transcript number - dd_Smed_v6_9413_0_1. For other species, Pfam accession numbers are as follows: human - Q9BSF4, mouse - Q8BGX2, *D. melanogaster* - Q9W4R8, *C. elegans* - Q8WQD7, tardigrade - A0A1D1VF99, octopus - A0A0L8FIM2, *N. vectensis* - A7SPD9, *C. intestinalis* - H2XT55, sponge - A0A1×7SN56.

**Supplementary Table 3.** Enrichment of neoblast and germline transcripts in differentially expressed genes in various comparisons.

| Dataset | Expresssion Pattern | Germline | Neoblasts |
| --- | --- | --- | --- |
| *DUF2366*(RNAi) Regeneration compared to *DUF2366*(RNAi) Homeostasis | Up | 0.02 | 0.44 |
|  | Down | 5.01 | 1.18 |
| *gfp*(RNAi) Regeneration compared to *gfp*(RNAi) Homeostasis | Up | 0.02 | 1.37 |
|  | Down | 5.58 | 0.40 |
| *DUF2366*(RNAi) Regeneration compared to *gfp*(RNAi) Regeneration | Up | 0.09 | 0.25 |
|  | Down | 0.68 | 4.77 |
| *DUF2366*(RNAi) Homeostatis compared to *gfp*(RNAi) Homeostasis | Up | ns | ns |
|  | Down | ns | ns |

**Supplementary Table 4.**
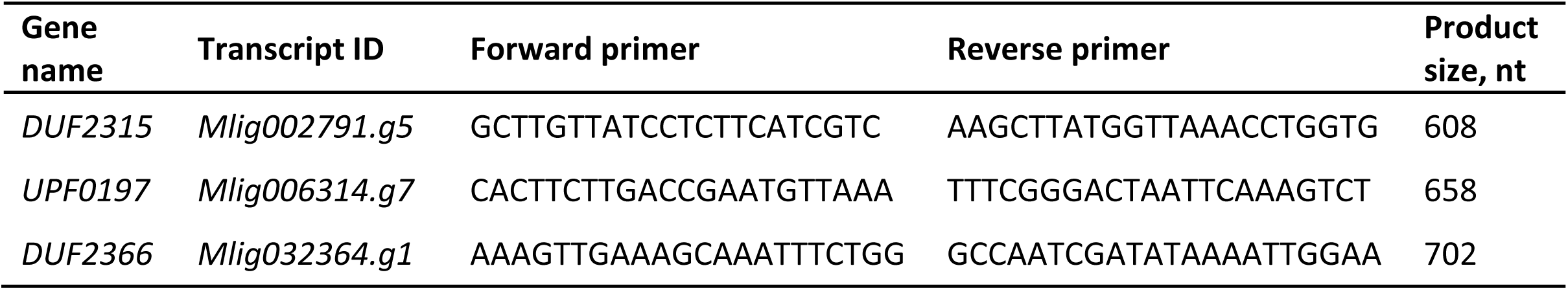
Primers for the production of dsRNA

## References

Anderson RA, Berges RA, Harrison PJ, Watanabe MM. 2005. Recipes for Freshwater and Seawater Media; Enriched Natural Seawater MediaAlgal Culturing Techniques. p. 596.

Bateman A, Coggill P, Finn RD. 2010. DUFs: families in search of function. Acta Crystallogr Sect F Struct Biol Cryst Commun 66:1148–1152. doi: 10.1107/S1744309110001685

Battersby BJ, Richter U. 2013. Why translation counts for mitochondria - retrograde signalling links mitochondrial protein synthesis to mitochondrial biogenesis and cell proliferation. J Cell Sci 126:4331–4338. doi: 10.1242/jcs.131888

Bode A, Salvenmoser W, Nimeth K, Mahlknecht M, Adamski Z, Rieger RM, Peter R, Ladurner P. 2006. Immunogold-labeled S-phase neoblasts, total neoblast number, their distribution, and evidence for arrested neoblasts in Macrostomum lignano (Platyhelminthes, Rhabditophora). Cell Tissue Res 325:577–587.

Brandl H, Moon HK, Vila-Farré M, Liu SY, Henry I, Rink JC. 2016. PlanMine - A mineable resource of planarian biology and biodiversity. Nucleic Acids Res 44:D764–D773. doi: 10.1093/nar/gkv1148

Callegari S, Richter F, Chojnacka K, Jans DC, Lorenzi I, Pacheu-Grau D, Jakobs S, Lenz C, Urlaub H, Dudek J, Chacinska A, Rehling P. 2016. TIM29 is a subunit of the human carrier translocase required for protein transport. FEBS Lett 590:4147–4158. doi: 10.1002/1873-3468.12450

Carlson M. 2019. org.Hs.eg.db: Genome wide annotation for Human. R Packag version 382.

Davidson NM, Oshlack A. 2014. Corset: enabling differential gene expression analysis for de novo assembled transcriptomes. Genome Biol 15:410. doi: 10.1186/s13059-014-0410-6

De Mulder K, Pfister D, Kuales G, Egger B, Salvenmoser W, Willems M, Steger J, Fauster K, Micura R, Borgonie G, Ladurner P. 2009. Stem cells are differentially regulated during development, regeneration and homeostasis in flatworms. Dev Biol 334:198–212. doi: 10.1016/j.ydbio.2009.07.019

Dobin A, Davis CA, Schlesinger F, Drenkow J, Zaleski C, Jha S, Batut P, Chaisson M, Gingeras TR. 2013. STAR: Ultrafast universal RNA-seq aligner. Bioinformatics 29:15–21. doi: 10.1093/bioinformatics/bts635

Durieux J, Wolff S, Dillin A. 2011. The Cell-Non-Autonomous Nature of Electron Transport Chain-Mediated Longevity. Cell 144:79–91. doi: 10.1016/j.cell.2010.12.016

Egger B, Ladurner P, Nimeth K, Gschwentner R, Rieger R. 2006. The regeneration capacity of the flatworm Macrostomum lignano - On repeated regeneration, rejuvenation, and the minimal size needed for regeneration. Dev Genes Evol 216:565–577. doi: 10.1007/s00427-006-0069-4

Fukasawa Y, Tsuji J, Fu S-C, Tomii K, Horton P, Imai K. 2015. MitoFates: improved prediction of mitochondrial targeting sequences and their cleavage sites. Mol Cell Proteomics 14:1113–26. doi: 10.1074/mcp.M114.043083

Goodacre NF, Gerloff DL, Uetz P. 2013. Protein domains of unknown function are essential in bacteria. MBio 5:e00744–13. doi: 10.1128/mBio.00744-13

Grudniewska M, Mouton S, Grelling M, Wolters AHG, Kuipers J, Giepmans BNG, Berezikov E. 2018. A novel flatworm-specific gene implicated in reproduction in Macrostomum lignano. Sci Rep 8:3192. doi: 10.1038/s41598-018-21107-4

Grudniewska M, Mouton S, Simanov D, Beltman F, Grelling M, de Mulder K, Arindrarto W, Weissert PM, van der Elst S, Berezikov E. 2016. Transcriptional signatures of somatic neoblasts and germline cells in *Macrostomum lignano*. Elife 5. doi: 10.7554/eLife.20607

Hashimshony T, Senderovich N, Avital G, Klochendler A, de Leeuw Y, Anavy L, Gennert D, Li S, Livak KJ, Rozenblatt-Rosen O, Dor Y, Regev A, Yanai I. 2016. CEL-Seq2: sensitive highly-multiplexed single-cell RNA-Seq. Genome Biol 17:77. doi: 10.1186/s13059-016-0938-8

Hashimshony T, Wagner F, Sher N, Yanai I. 2012. CEL-Seq: Single-Cell RNA-Seq by Multiplexed Linear Amplification. Cell Rep 2:666–673. doi: 10.1016/j.celrep.2012.08.003

Houtkooper RH, Mouchiroud L, Ryu D, Moullan N, Katsyuba E, Knott G, Williams RW, Auwerx J. 2013. Mitonuclear protein imbalance as a conserved longevity mechanism. Nature 497:451–457. doi: 10.1038/nature12188

Jaroszewski L, Li Z, Krishna SS, Bakolitsa C, Wooley J, Deacon AM, Wilson IA, Godzik A. 2009. Exploration of Uncharted Regions of the Protein Universe. PLoS Biol 7:e1000205. doi: 10.1371/journal.pbio.1000205

Jensen MB, Jasper H. 2014. Mitochondrial Proteostasis in the Control of Aging and Longevity. Cell Metab 20:214–225. doi: 10.1016/j.cmet.2014.05.006

Kang Y, Baker MJ, Liem M, Louber J, McKenzie M, Atukorala I, Ang CS, Keerthikumar S, Mathivanan S, Stojanovski D. 2016. Tim29 is a novel subunit of the human TIM22 translocase and is involved in complex assembly and stability. Elife 5:1–22. doi: 10.7554/eLife.17463

Ladurner P, Egger B, Mulder K De, Pfister D, Kuales G, Salvenmoser W, Schärer L. 2008. The Stem Cell System of the Basal Flatworm Macrostomum lignanoStem Cells: From Hydra to Man. Berlin - Heidelberg - New York: Bosh, Th.C.G. pp. 75–94.

Ladurner P, Rieger R, Baguñà J. 2000. Spatial distribution and differentiation potential of stem cells in hatchlings and adults in the marine platyhelminth *macrostomum sp.*: a bromodeoxyuridine analysis. Dev Biol 226:231–241. doi: 10.1006/dbio.2000.9867

Lin YF, Haynes CM. 2016. Metabolism and the UPRmt. Mol Cell. doi: 10.1016/j.molcel.2016.02.004

Lisowski P, Kannan P, Mlody B, Prigione A. 2018. Mitochondria and the dynamic control of stem cell homeostasis. EMBO Rep 19:e45432. doi: 10.15252/embr.201745432

Lu S, Wang J, Chitsaz F, Derbyshire MK, Geer RC, Gonzales NR, Gwadz M, Hurwitz DI, Marchler GH, Song JS, Thanki N, Yamashita RA, Yang M, Zhang D, Zheng C, Lanczycki CJ, Marchler-Bauer A. 2020. CDD/SPARCLE: the conserved domain database in 2020. Nucleic Acids Res 48:D265–D268. doi: 10.1093/nar/gkz991

McCarthy DJ, Chen Y, Smyth GK. 2012. Differential expression analysis of multifactor RNA-Seq experiments with respect to biological variation. Nucleic Acids Res 40:4288–4297. doi: 10.1093/nar/gks042

Mouton S, Wudarski J, Grudniewska M, Berezikov E. 2018. The regenerative flatworm Macrostomum lignano, a model organism with high experimental potential. Int J Dev Biol 62:551–558. doi: 10.1387/ijdb.180077eb

Mudgal R, Sandhya S, Chandra N, Srinivasan N. 2015. De-DUFing the DUFs: Deciphering distant evolutionary relationships of Domains of Unknown Function using sensitive homology detection methods. Biol Direct 10:38. doi: 10.1186/s13062-015-0069-2

Nimeth K, Ladurner P, Gschwentner R, Salvenmoser W, Rieger R. 2002. Cell renewal and apoptosis in Macrostomum sp. [Lignano]. Cell Biol Int 26:801–815. doi: 10.1016/S1065-6995(02)90950-9

Nimeth KT, Egger B, Rieger R, Salvenmoser W, Peter R, Gschwentner R. 2007. Regeneration in Macrostomum lignano (Platyhelminthes): cellular dynamics in the neoblast stem cell system. Cell Tissue Res 327:637–46. doi: 10.1007/s00441-006-0299-9

Okonechnikov K, Golosova O, Fursov M. 2012. Unipro UGENE: a unified bioinformatics toolkit. Bioinformatics 28:1166–1167. doi: 10.1093/bioinformatics/bts091

Pfister D, De Mulder K, Hartenstein V, Kuales G, Borgonie G, Marx F, Morris J, Ladurner P. 2008. Flatworm stem cells and the germ line: Developmental and evolutionary implications of macvasa expression in Macrostomum lignano. Dev Biol 319:146–159. doi: 10.1016/j.ydbio.2008.02.045

Pfister D, De Mulder K, Philipp I, Kuales G, Hrouda M, Eichberger P, Borgonie G, Hartenstein V, Ladurner P, Egger B, Obwegeser S, Salvenmoser W, Ladurner P. 2007. The exceptional stem cell system of Macrostomum lignano: Screening for gene expression and studying cell proliferation by hydroxyurea treatment and irradiation. Front Zool 4:9. doi: 10.1186/1742-9994-4-9

Rafalski VA, Mancini E, Brunet A. n.d. Energy metabolism and energy-sensing pathways in mammalian embryonic and adult stem cell fate. J Cell Sci 125:5597–5608. doi: 10.1242/jcs.114827

Rieger R, Gehlen M, Haszprunar G, Holmlund M, Legniti A, Salvenmoser W, Tyler S. 1988. Laboratory cultures of marine Macrostomida (Turbellaria). Fortschr Zool 36.

Rieger RM, Legniti A, Ladurner P, Reiter D, Asch E, Salvenmoser WS, SchÜrmann W, Peter R. 1999. Ultrastructure of neoblasts in microturbellaria: Significance for understanding stem cells in free-living platyhelminthes. Invertebr Reprod Dev 35:127–140. doi: 10.1080/07924259.1999.9652376

Samluk L, Chroscicki P, Chacinska A. 2018. Mitochondrial protein import stress and signaling. Curr Opin Physiol 3:41–48. doi: 10.1016/j.cophys.2018.02.010

Wilkening S, Pelechano V, Järvelin AI, Tekkedil MM, Anders S, Benes V, Steinmetz LM. 2013. An efficient method for genome-wide polyadenylation site mapping and RNA quantification. Nucleic Acids Res 41:6370–6370. doi: 10.1093/nar/gkt364

Wudarski J, Egger B, Ramm SA, Schärer L, Ladurner P, Zadesenets KS, Rubtsov NB, Mouton S, Berezikov E. 2020. The free-living flatworm Macrostomum lignano. Evodevo 11:5. doi: 10.1186/s13227-020-00150-1

Wudarski J, Simanov D, Ustyantsev K, de Mulder K, Grelling M, Grudniewska M, Beltman F, Glazenburg L, Demircan T, Wunderer J, Qi W, Vizoso DB, Weissert PM, Olivieri D, Mouton S, Guryev V, Aboobaker A, Schärer L, Ladurner P, Berezikov E. 2017. Efficient transgenesis and annotated genome sequence of the regenerative flatworm model Macrostomum lignano. Nat Commun 8:2120. doi: 10.1038/s41467-017-02214-8

Yoneda T, Benedetti C, Urano F, Clark SG, Harding HP, Ron D. 2004. Compartment-specific perturbation of protein handling activates genes encoding mitochondrial chaperones. J Cell Sci 117:4055–4066. doi: 10.1242/jcs.01275

Yu G, Wang L-G, Han Y, He Q-Y. 2012. clusterProfiler: an R Package for Comparing Biological Themes Among Gene Clusters. Omi A J Integr Biol 16:284–287. doi: 10.1089/omi.2011.0118

Zhang H, Menzies KJ, Auwerx J. 2018. The role of mitochondria in stem cell fate and aging. Development 145:dev143420. doi: 10.1242/dev.143420

